# Limited effects of age on the use of the ankle and counter-rotation mechanism in the sagittal plane

**DOI:** 10.1101/2022.03.15.484389

**Authors:** Maud van den Bogaart, Sjoerd M. Bruijn, Joke Spildooren, Jaap H. van Dieën, Pieter Meyns

## Abstract

Two mechanisms can be used to accelerate the center of mass (CoM) to control the CoM in relation to the base of support during standing. The first is applying ankle moments to shift the center of pressure (CoP), which has been coined the “CoP mechanism”. The second is changing the angular momentum around the CoM to change the direction of the ground reaction force, i.e., the “counter-rotation mechanism”. At both the beginning and the end of the lifespan, problems with postural control are common. In this study, we asessed anteroposterior balance performance and the related use of these postural control mechanisms in children, younger adults, and older adults. Sixteen pre-pubertal children (6-9y), 17 younger adults (18-24y) and eight older adults (65-80y) performed bipedal upright standing trials of 16 seconds on a rigid surface and on three balance boards that could freely move in the sagittal plane, varying in height (15-19 cm) of the board above the point of contact with the floor. Full body kinematics were measured. Performance related outcome measures, i.e., the number of trials with balance loss and the Root Mean Square (RMS) of the time series of the CoM acceleration were calculated. Additionally, the RMS of the time series of the CoM acceleration due to the CoP and counter-rotation mechanism and the contributions of the CoP and the counter-rotation mechanism to the CoM acceleration (in %) in the sagittal plane were calculated. Furthermore, selected kinematic measures, i.e., the orientation of the board and the head and the Mean Power Frequency of balance board orientation and of CoM acceleration were calculated. Compared to younger adults, children and older adults showed a poorer balance performance, reflected by a greater RMS of CoM accelerations and more balance loss in older adults. Across age groups and conditions, the contribution of the CoP mechanism to the total CoM acceleration was dominant, i.e., 95%-108%. The contribution of the counter-rotation mechanism ranged between 19%-31% (with totals higher than 100% indicating opposite effects of both mechanisms). We suggest that the contribution of the counter-rotation mechanism is limited, since the counter-rotation mechanism would conflict with stabilizing the orientation of the head in space. Furthermore, children did use the counter-rotation mechanism relatively more to accelerate the CoM compared to younger adults. Possibly this reflects that they are still learning to limit the contribution of the counter-rotation mechanism to the same extent as adults.

## 1. Introduction

Adequate postural control is a prerequisite for performance of crucial activities of daily life. Postural control is necessary to prevent falls and can be defined as controlling the state of the body center of mass (CoM, i.e., the point around which the mass is evenly distributed) relative to the base of support (BoS, i.e., the area within an outline of all points on the body which are directly in contact with the support surface) (Horak, 1987). Postural control is regulated by the sensorimotor control system; this integrates sensory input from visual, vestibular and somatosensory systems to generate motor commands, resulting in muscular responses or motor output (Peterka, 2002).

During standing, two postural control mechanisms can be used to accelerate the CoM to control the CoM in relation to the base of support during standing. The first of these is activating muscles around the ankle to generate ankle moments (Hof, 2007; Horak et al., 1986). These ankle moments are reflected in a shift of the center of pressure of the ground reaction force (CoP). Consequently, this mechanism has been coined the “CoP mechanism” (Hof, 2007; Horak et al., 1986). The second mechanism is changing the angular momentum around the CoM to change the direction of the ground reaction force, i.e., the “counter-rotation mechanism” (Hof, 2007). Rotation of the trunk and pelvis around the hip, which has been called the hip mechanism in the literature, is one example of the counter-rotation mechanism (Hof, 2007; Otten, 1999). Other examples of the counter-rotation mechanism are accelerations of other body segments, such as the arms or head, which can be used in the same way. The use of these postural control mechanisms has been suggested to be direction-specific (Winter et al., 1998; Winter et al., 1996). Therefore, in a previous paper, we focused on the use of the postural control mechanism in the frontal plane (van den Bogaart et al., 2022). The current manuscript focusses on the use of the postural control mechanisms in the sagittal plane in the same population.

During quiet bipedal stance, the ankle mechanism is the dominant mechanism to accelerate the CoM in the sagittal plane in healthy younger adults (Winter et al., 1996). More proximal muscles will be activated when standing on a compliant or moving support surface (Patel et al., 2008; Riemann et al., 2003). Standing on such a surface makes proprioceptive information at the ankle less pertinent and changes the effects of ankle moments on CoM acceleration (Horak et al., 2001; MacLellan et al., 2006). Therefore, it is expected that people will rely more on the counter-rotation mechanism with increasing surface instability. The frequency of postural corrections made using the CoP mechanism could be increased when standing on a moving support surface as the mean power frequency (MPF) of CoP displacements is higher when standing on a sway-referenced support surface compared to a rigid surface (Dickin et al., 2012).

At both the beginning and the end of the lifespan, problems with postural control are common. In children, immature sensory systems limit postural control (Steindl et al., 2006). Maturation of the somatosensory system occurs at 3 to 4 years of age and the visual and vestibular systems reach adult levels at 15 to 16 years of age (Steindl et al., 2006) or even later (Hirabayashi et al., 1995). The integration and reweighting of sensory information is not yet adult-like until the age of 15 (Shams et al., 2020). This could explain the differences in balance performance between children and adults. During quiet standing and standing on foam, the amplitude of CoP displacements, CoP velocities and CoM accelerations and the MPF of CoM accelerations, are larger in children between 3 and 6 years old than in older children and adults (Hsu et al., 2009; Oba et al., 2015). In addition to the differences in balance performance, children between 4 and 6 years old showed variable use of postural control mechanisms after disturbances of upright standing by a movable platform. Sometimes the children demonstrated an ankle mechanism, and sometimes they demonstrated a counter-rotation mechanism (specifically the hip mechanism) (Shumway-Cook et al., 1985). It was postulated that children do not show adult-like consistent use of the postural control mechanisms until 10 years of age (Shumway-Cook et al., 1985). Information on the use of the CoP mechanism and counter-rotation mechanism when standing on different (unstable) surfaces in children is, to the best of our knowledge, missing.

In older adults, deterioration of the sensory and motor systems, as well as sensory reweighting deficits occur (Sturnieks et al., 2008). Deficits in the sensorimotor control system in older adults lead to impaired balance performance compared to younger adults (Toledo et al., 2010). CoM accelerations and MPF of CoP velocities were larger in older adults (age > 70) compared to younger adults during quiet standing (Demura et al., 2006; Masani et al., 2007; Yu et al., 2008). The amplitude of CoP displacements was larger after forward platform translations when comparing older adults (age > 65) with younger adults (Nakamura et al., 2001). When comparing the use of the postural control mechanisms between older and young people, older adults tend to use the counter-rotation mechanism more after perturbations of standing (Gu et al., 1996; Liaw et al., 2009; Lin et al., 2004; Manchester et al., 1989). Information on the use of the CoP mechanism and counter-rotation mechanism in the AP direction when standing on different (unstable) surfaces in older adults is still missing and worthwhile to assess.

We assessed if, and how, children, younger adults and older adults use the counter-rotation mechanism to accelerate the CoM during standing and how this interacts with the CoP mechanism, during standing on unstable support surfaces, i.e., uniaxial balance boards that can freely move in the sagittal plane. To test if, and how, balance performance and the related use of the postural control mechanisms change with ageing, variations in surface instability were used. We expected poorer balance performance and more use of the counter-rotation mechanism in children and older adults compared to younger adults. We also expected poorer balance performance and increased use of the counter-rotation mechanism during standing on the balance boards compared to standing on a rigid surface. Additionally, we hypothesized that the CoP mechanism is dominant, based on our findings when assessing the use of the postural control mechanisms in the frontal plane (van den Bogaart et al., 2022).

## 2. Methods

The methods and participants of this study were identical to that of our previous study (van den Bogaart et al., 2022). In the current experimental setup, however, the direction of the movement of the balance boards was in the anteroposterior direction (i.e., in the sagittal plane). Whereas in our previous study the balance boards only allowed movement in the mediolateral direction (i.e., in the frontal plane).

### 2.1. Subjects

Sixteen pre-pubertal children between 6-9 years old (10 males, age 8.2±1.1 years old, BMI 15.6±1.5 kg/m^2^), 17 healthy younger adults between 18-24 years old (7 males, age 21.9±1.6 years old, BMI 23.5±3.0 kg/m^2^) and eight older adults between 65-80 years old (5 males, age 71.8±4.6 years old, BMI 26.0±3.4 kg/m^2^) participated. The required sample size calculated was eight per group (van den Bogaart et al., 2022). Participants gave written informed consent prior to the experiment. The study protocol was in agreement with the declaration of Helsinki and had been approved by the local ethical committee (CME2018/064, NCT04050774).

### 2.2. Research design

The participants performed bipedal upright standing on a rigid surface and on three balance boards varying in height of the surface above the point of contact with the floor (BB1; 15 cm, BB2; 17 cm and BB3; 19 cm). The balance board was a 48 cm by 48 cm wooden board mounted on a section of a cylinder with a 24 cm radius that could freely move in the sagittal plane (Figure 1). The four conditions were repeated three times in random order, with each trial lasting 16 seconds. For every trial, participants were instructed to stand barefoot on two feet, placed in parallel at hip width and arms along the body. They were asked to look at a marked spot at seven meters distance on the wall in front of them at eye level.

**Figure 1.**
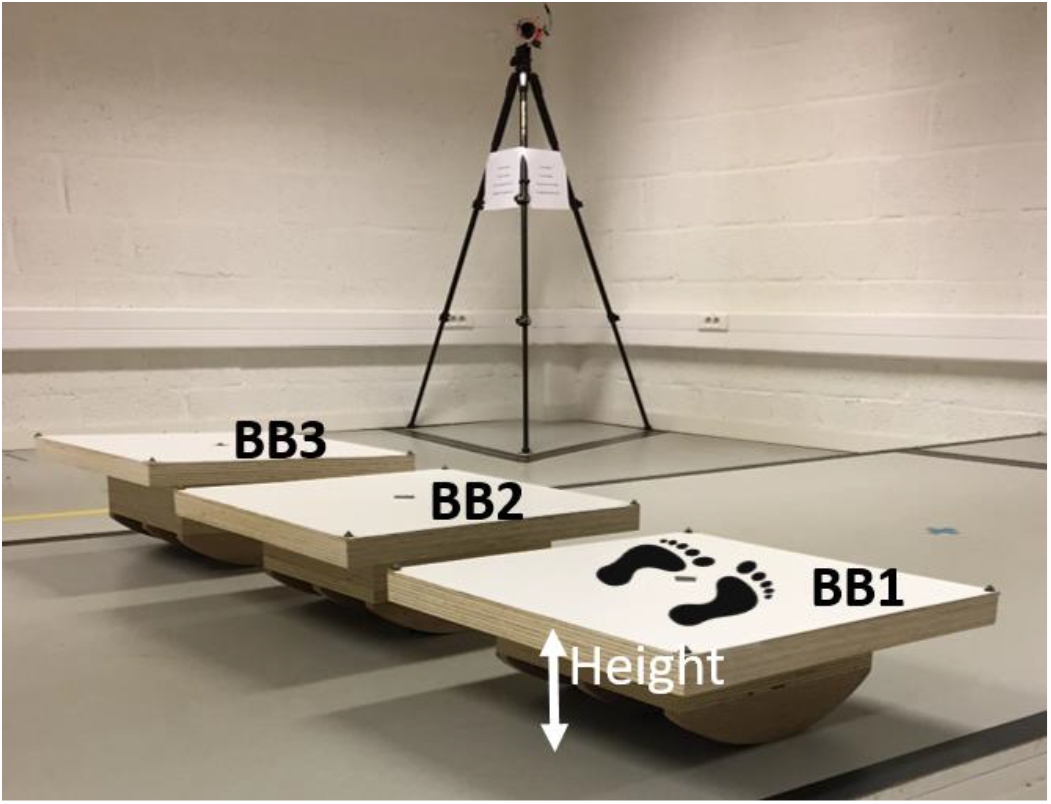
Illustration of the balance boards, which could freely move in the sagittal plane, varying in height of the surface of the board above the point of contact with the floor (BB1; 15 cm, BB2; 17 cm and BB3; 19 cm). The feet indicate the person’ orientation on the balance board.

### 2.3. Materials and software

A Simi 3D motion analysis system (GmbH) with eight cameras (sample rate: 100 samples/sec, resolution: 1152×864 pixels) and 48 retro reflective markers was used. The illumination in the room at eye level was 650 Lux. Full body 3D kinematics (16 segments) were retrieved using the open-source deep learning python toolboxes DeepLabCut (https://github.com/AlexEMG/DeepLabCut) and Anipose (https://github.com/lambdaloop/anipose). The complete workflow has been described previously (van den Bogaart et al., 2022).

### 2.4 Data analysis

#### 2.4.1. Performance

A trial was registered as a balance loss if a stepping response or an intervention by a researcher was required in order to remain standing. The number of balance losses per condition and per age group was recorded as a performance related outcome measure. In case of balance loss, the trial was excluded from further analysis. Next to the number of balance losses, the Root Mean Square (RMS) of the time series of the CoM acceleration in the sagittal plane was determined as a measure of performance.

#### 2.4.2. Postural control mechanisms

The magnitudes of CoM acceleration induced by the CoP mechanism and counter-rotation mechanism in the sagittal plane were calculated using Eq. (1), as described by Hof (2007).

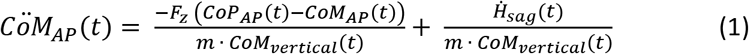

in which *m* is body mass, *CoM_AP_* is the anteroposterior (AP) position of the CoM, *CoM_vertical_* is the vertical position of the CoM, *CÖM_AP_* is the double derivative of *CoM_AP_* with respect to time, *t* is time, *F_z_* is the vertical ground reaction force, *CoP_AP_* is the AP position of the CoP, and 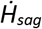 is the change in total body angular momentum in the sagittal plane.

Here, the first part of the right-hand term, 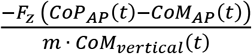, refers to the CoP mechanism and the second part, 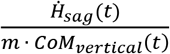, is the AP CoM acceleration induced by the counter-rotation mechanism.

As it was not possible to collect accurate ground reaction forces and CoP, the magnitude of AP CoM acceleration induced by the CoP mechanism was calculated by subtracting, 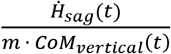, from *CÖM_Ap_*(*t*).

The RMS values of the time series of CoM acceleration due to CoP and counter-rotation mechanism were calculated for each trial. The relative contributions of the CoP and counter-rotation mechanism to the *CöM_AP_* (in %) were calculated by dividing the RMS of each mechanism by *CöM_AP_*, multiplied by 100. Totals higher than 100% indicate opposite effects of both mechanisms.

#### 2.4.3. Kinematics

Orientations of the board and head in the sagittal plane were calculated relative to the global coordinate system. The deviations from the mean orientations and the MPF of the balance board orientation and *CöM_AP_* were calculated as described previously (van den Bogaart et al., 2022).

### 2.5. Statistics

The number of trials for each surface condition was three unless balance loss occurred, which resulted in exclusion of this trial. Fisher exact tests were used to compare the number of balance losses of older adults and children with younger adults. The results of the successful trials of each surface condition were averaged for each participant. Two-way repeated measures ANOVAs were used to determine the effect of Age and Surface as well as their interaction on the RMS of *CöM_AP_*, RMS of CoM acceleration due to the CoP and counter-rotation mechanism, the relative contribution of the CoP and counter-rotation mechanism to *CöM_AP_*, the SD of the balance board and head orientation, and the MPF of *CöM_AP_* and balance board orientation. Post-hoc analyses were performed to determine differences between the surface conditions (using a Bonferroni correction of the p-values). In addition, planned comparisons (without correction for multiple testing) were done to compare children with younger adults and older adults with younger adults. Statistical analyses were performed with SPSS(v25) with α<0.05.

## 3. Results

In spite of slight deviations from normality, parametric statistical testing was performed. Transforming data hampers the interpretation of the results and ANOVA is considered robust to violations of normality (Schmider et al., 2010).

### 3.1. Performance

#### 3.1.1. Balance loss

None of the participants had to be excluded because at least one out of the three trials per surface condition per participant was available. Older adults did lose balance more often than younger adults, 50% versus 5.9% respectively (Table 1, *p* = 0.023).

**Table 1.** The number of balance losses per surface condition (standing on a rigid surface (RIGID) and uniaxial balance boards varying in height BB1; 15 cm, BB2; 17 cm and BB3; 19 cm).

|  | Surface condition |  |  |  |
| --- | --- | --- | --- | --- |
|  | RIGID | BB1 | BB2 | BB3 |
| Child |  | 1x |  |  |
| Child |  | 1x |  |  |
| Child |  | 1x | 1x |  |
| Younger adult |  |  |  | 1x |
| Older adult |  | 2x | 2x | 2x |
| Older adult |  | 1x | 1x | 2x |
| Older adult |  | 2x | 1x | 1x |
| Older adult |  | 1x | 2x | 1x |

#### 3.1.2 Total CoM acceleration

Significant main effects of Age, Surface and a significant interaction of Age and Surface on the RMS of *CöM_AP_* were found (Table 2, Figure 2a). The RMS of *CöM_AP_*was significantly larger in children and older adults compared to younger adults across all conditions. In addition, the RMS of *CöM_AP_* was significantly smaller during standing on a rigid surface compared to standing on the balance boards across all age groups. The RMS of *CöM_AP_* was significantly smaller during standing on a rigid surface compared to standing on BB1 in children and younger adults, but not significantly different in older adults (*p* = 0.052).

**Figure 2.**
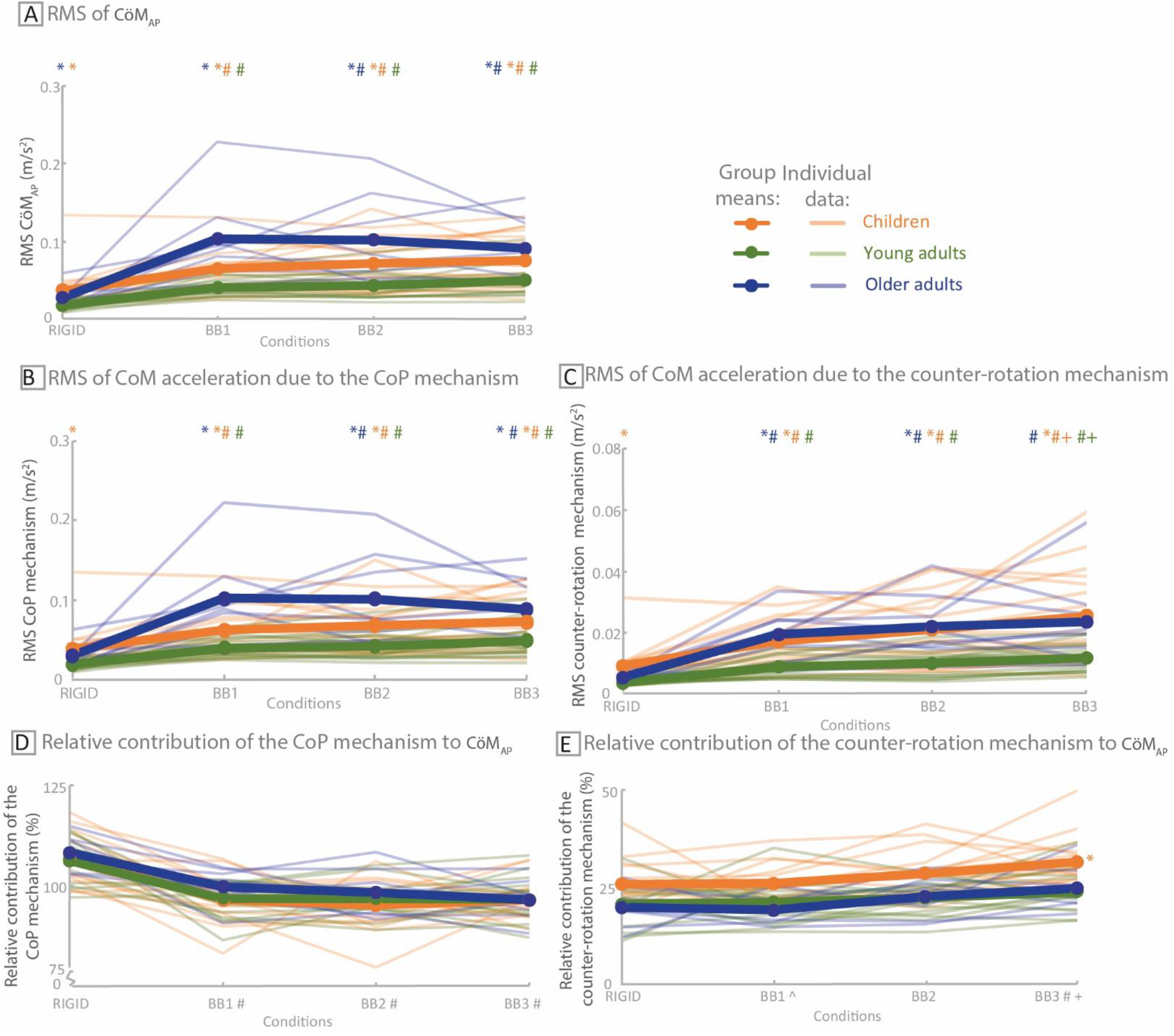
Group means (thick lines) and individual data (thin lines) of the **A)** Root Mean Square (RMS) of the Center of Mass (CoM) acceleration (*CöM_AP_*) (in m/s^2^), **B)** the RMS of CoM acceleration due to the CoP mechanism (in m/s^2^), **C)** the RMS of CoM acceleration due to the counter-rotation mechanism (including zooming in) (in m/s^2^), **D)** the relative contribution of the CoP mechanism to *CöM_AP_* (in %), **E)** the relative contribution of the counter-rotation mechanism to *CöM_AP_* (in %), during standing on a rigid surface (RIGID) and during standing on uniaxial balance boards that can freely move in anteroposterior direction varying in height (BB1; 15 cm, BB2; 17 cm and BB3; 19 cm) in children (orange), younger adults (green) and older adults (blue). * represents a significant difference compared to younger adults, with the group tested identified by the color code. # represents a significant difference compared to standing on a rigid surface. + represents a significant difference compared to standing on BB1. ^ represents a significant difference compared to standing on BB2.

**Table 2.**
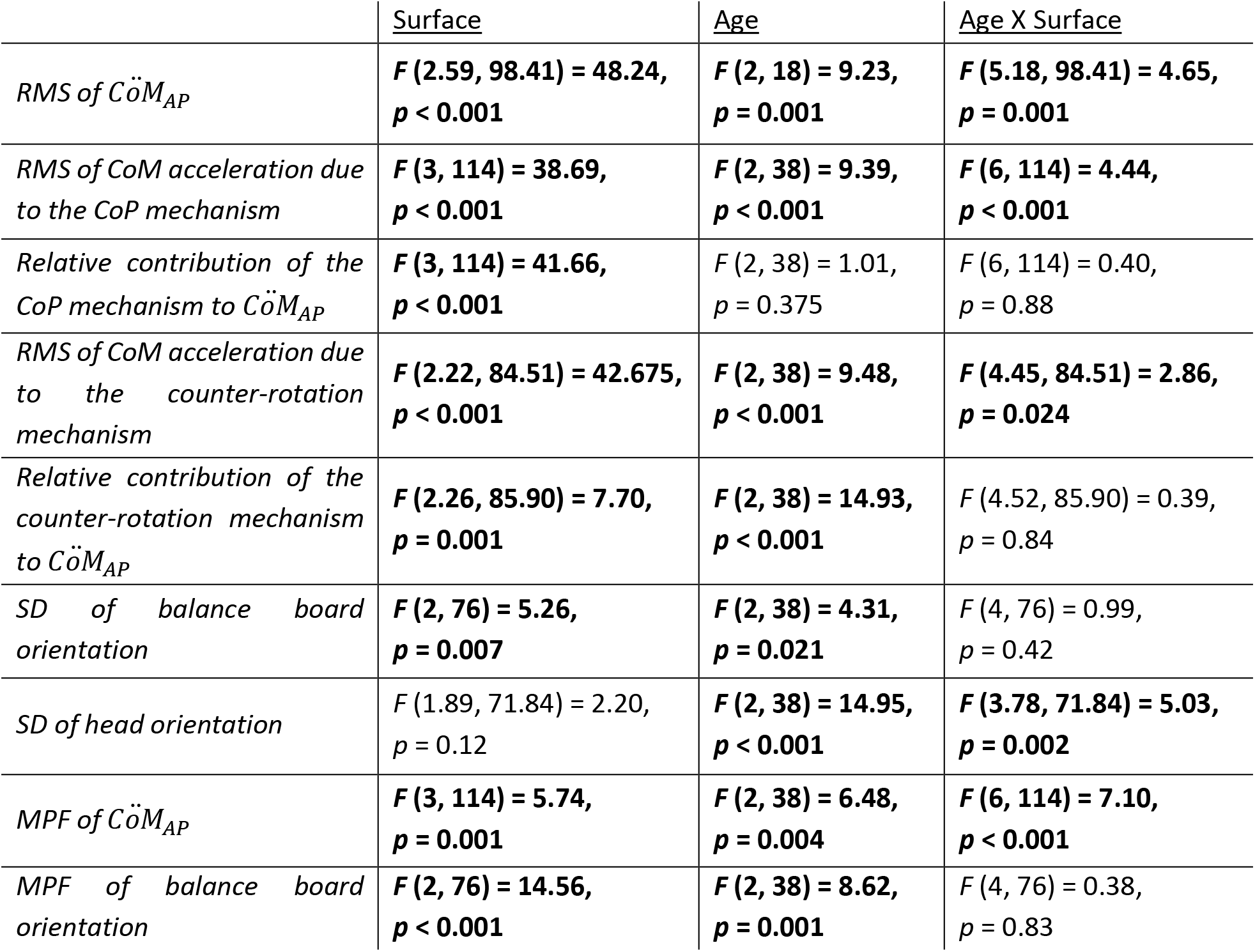
Two-way repeated measures ANOVA results for the effect of Age and Surface as well as their interaction on dependent variables. Bold text corresponds to significant effects.

|  | <u>Surface</u> | <u>Age</u> | <u>Age X Surface</u> |
| --- | --- | --- | --- |
| *RMS of $\ddot{C}oM_{AP}$* | **$F(2.59, 98.41) = 48.24$,**<br>**$p < 0.001$** | **$F(2, 18) = 9.23$,**<br>**$p = 0.001$** | **$F(5.18, 98.41) = 4.65$,**<br>**$p = 0.001$** |
| *RMS of CoM acceleration due to the CoP mechanism* | **$F(3, 114) = 38.69$,**<br>**$p < 0.001$** | **$F(2, 38) = 9.39$,**<br>**$p < 0.001$** | **$F(6, 114) = 4.44$,**<br>**$p < 0.001$** |
| *Relative contribution of the CoP mechanism to $\ddot{C}oM_{AP}$* | **$F(3, 114) = 41.66$,**<br>**$p < 0.001$** | $F(2, 38) = 1.01$,<br>$p = 0.375$ | $F(6, 114) = 0.40$,<br>$p = 0.88$ |
| *RMS of CoM acceleration due to the counter-rotation mechanism* | **$F(2.22, 84.51) = 42.675$,**<br>**$p < 0.001$** | **$F(2, 38) = 9.48$,**<br>**$p < 0.001$** | **$F(4.45, 84.51) = 2.86$,**<br>**$p = 0.024$** |
| *Relative contribution of the counter-rotation mechanism to $\ddot{C}oM_{AP}$* | **$F(2.26, 85.90) = 7.70$,**<br>**$p = 0.001$** | **$F(2, 38) = 14.93$,**<br>**$p < 0.001$** | $F(4.52, 85.90) = 0.39$,<br>$p = 0.84$ |
| *SD of balance board orientation* | **$F(2, 76) = 5.26$,**<br>**$p = 0.007$** | **$F(2, 38) = 4.31$,**<br>**$p = 0.021$** | $F(4, 76) = 0.99$,<br>$p = 0.42$ |
| *SD of head orientation* | $F(1.89, 71.84) = 2.20$,<br>$p = 0.12$ | **$F(2, 38) = 14.95$,**<br>**$p < 0.001$** | **$F(3.78, 71.84) = 5.03$,**<br>**$p = 0.002$** |
| *MPF of $\ddot{C}oM_{AP}$* | **$F(3, 114) = 5.74$,**<br>**$p = 0.001$** | **$F(2, 38) = 6.48$,**<br>**$p = 0.004$** | **$F(6, 114) = 7.10$,**<br>**$p < 0.001$** |
| *MPF of balance board orientation* | **$F(2, 76) = 14.56$,**<br>**$p < 0.001$** | **$F(2, 38) = 8.62$,**<br>**$p = 0.001$** | $F(4, 76) = 0.38$,<br>$p = 0.83$ |

### 3.2 Postural control mechanisms

#### 3.2.1. CoP mechanism

Significant main effects of Age, Surface and a significant interaction of Age and Surface on the RMS of the contribution of the CoP mechanism were found (Table 2, Figure 2b). The RMS of CoM acceleration due to the CoP mechanism was significantly larger in children and older adults compared to younger adults across all conditions. Furthermore, the RMS of CoM acceleration due to the CoP mechanism was significantly smaller during standing on a rigid surface compared to standing on the balance boards across all age groups. The RMS of CoM acceleration due to the CoP mechanism was significantly larger in older adults compared to younger adults in the balance board conditions, but no significant difference was found between younger adults and older adults when standing on a rigid surface (*p* = 0.056). Moreover, the RMS of CoM acceleration due to the CoP mechanism was significantly smaller during standing on a rigid surface compared to standing on BB1 in children and younger adults, but not in older adults (*p* = 0.097).

The relative contribution of the CoP mechanism to *CöM_AP_* ranged from 95%-108%. (Figure 2d). The average relative contribution decreased from standing on a rigid surface to standing on the balance boards (effect of Surface, Table 2 and Figure 2d).

#### 3.2.2. Counter-rotation mechanism

Significant main effects of Age, Surface and a significant interaction of Age and Surface on the RMS of CoM acceleration due to the counter-rotation mechanism were found (Table 2, Figure 2c). The RMS of CoM acceleration due to the counter-mechanism was significantly larger in children and older adults compared to younger adults across all conditions. Moreover, the RMS of CoM acceleration due to the counter-rotation mechanism was significantly smaller during standing on a rigid surface compared to standing on the balance boards across all groups. In addition, the RMS of CoM acceleration due to the counter-rotation mechanism was significantly smaller during standing on BB1 compared to standing on BB3 across all groups. The RMS of CoM acceleration due to the counter-mechanism was significantly larger in older adults compared to younger adults during standing on BB1 and BB2, but not when standing on a rigid surface (*p* = 0.052) and on BB3 (*p* = 0.058). The RMS of CoM acceleration due to the counter-rotation mechanism increased significantly with increasing height of the balance board in children and younger adults, with differences between standing on BB1 and BB3, but did not significantly increase in older adults.

The relative contribution of the counter-rotation mechanism to *CöM_AP_* ranged from 19%-31%. (Figure 2e). The relative contribution of the counter-rotation mechanism was significantly larger in children compared to younger adults (effect of Age, Table 2), but was not different between older and younger adults. Moreover, the relative contribution of the counter-rotation mechanism to *CöM_AP_* increased with surface instability, with differences between standing on a rigid surface and BB1 on one hand and BB3 on the other hand, and between standing on BB1 and BB2 (Effect of Surface).

### 3.3 Kinematics

#### 3.3.1. Balance board orientation

The SD of balance board orientation was larger in older adults compared to younger adults (effect of Age, Table 2 and Figure 3a). The SD of balance board orientation was significantly smaller when standing on BB1 than when standing on BB3 (effect of Surface).

**Figure 3.**
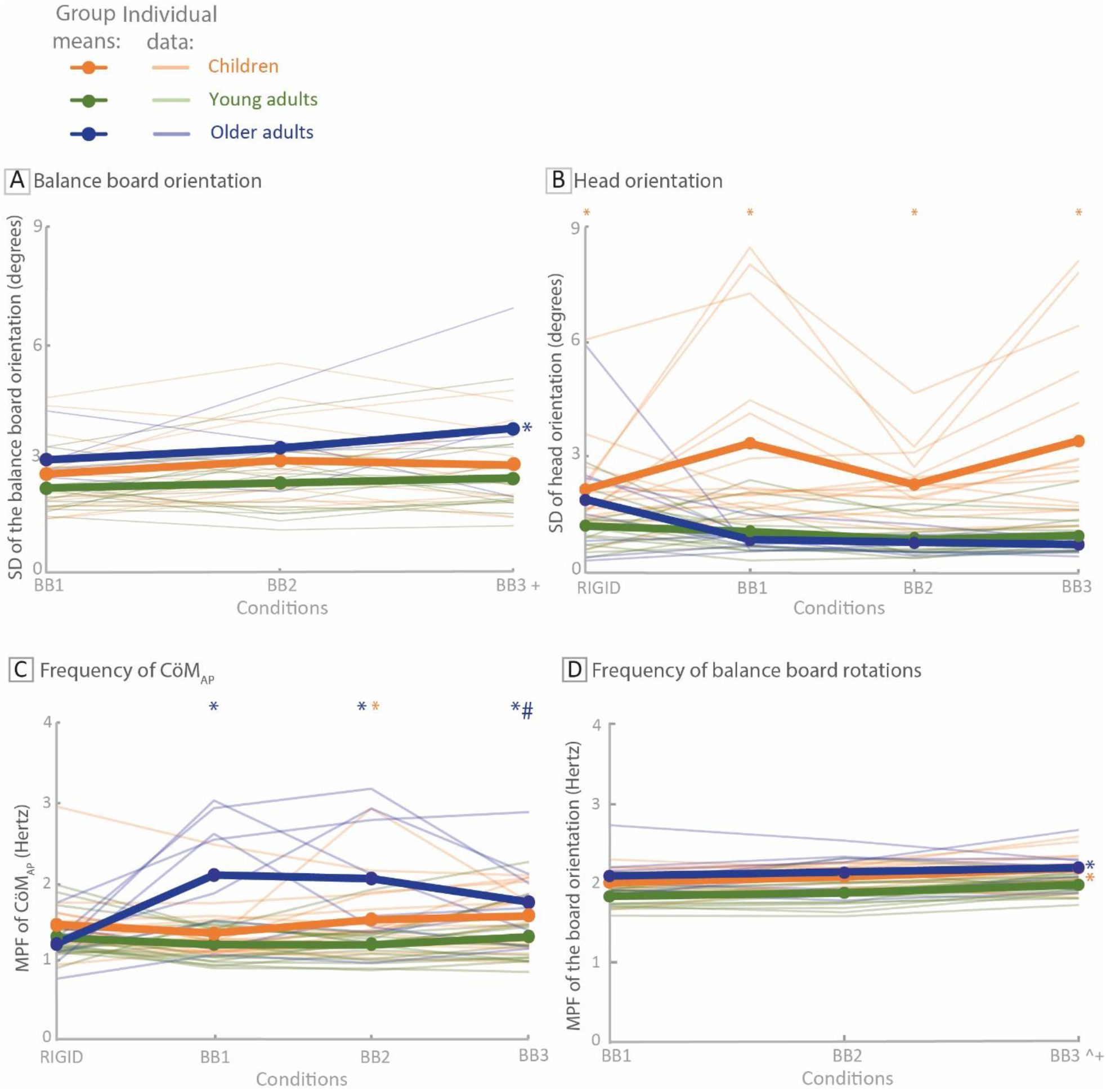
Group means (thick lines) and individual data (thin lines) of the **A)** standard deviation (SD) of the balance board orientation (in degrees), **B)** SD of the head orientation (in degrees), **C)** Mean Power Frequency (MPF) of *CöM_AP_* (in Hertz), **D)** MPF of the board orientation (in Hertz), during standing on a rigid surface (RIGID) and during standing on uniaxial balance boards that can freely move in anteroposterior direction varying in height (BB1; 15 cm, BB2; 17 cm and BB3; 19 cm) in children (orange), younger adults (green) and older adults (blue). * represents a significant difference compared to younger adults, with the group tested identified by the color code. # represents a significant difference compared to standing on a rigid surface. + represents a significant difference compared to standing on BB1. ^ represents a significant difference compared to standing on BB2.

#### 3.3.2. Head orientation

A significant main effect of Age and a significant interaction of Age and Surface on the SD of head orientation were found (Table 2, Figure 3b). The SD of head rotation was significantly larger in children compared to younger adults across all conditions. Post-hoc tests did not reveal significant effects.

#### 3.3.3. MPF of CoM accelerations and balance board rotations

Significant main effects of Age, Surface and a significant interaction of Age and Surface on the MPF of *CöM_AP_* were found (Table 2, Figure 3c). The MPF of *CöM_AP_* was significantly larger in older adults compared to younger adults across all conditions. Moreover, the MPF of *CöM_AP_* was significantly smaller during standing on a rigid surface compared to standing on the balance boards across all groups. The MPF of *CöM_AP_* was significantly larger in children compared to younger adults when standing on BB2 (BB3; *p* = 0.055). Furthermore, the MPF of *CöM_AP_* was significantly larger in older adults compared to younger adults during all balance board conditions. In older adults, the MPF of *CöM_AP_* was significantly lower during standing on the rigid surface compared to standing on BB3, but not so in younger adults and children.

The MPF of balance board orientation was significantly larger in children and older adults compared to younger adults (effect of Age, Table 2, Figure 3d). Moreover, the MPF of balance board orientation increased with surface instability, with differences between standing on BB1 and BB2 on one hand and standing on BB3 on the other hand (effect of Surface).

## 4. Discussion

We assessed if, and how, children, younger adults and older adults use the counter-rotation mechanism to accelerate their CoM during standing and how this interacts with the CoP mechanism, during standing on moving support surfaces, i.e., uniaxial balance boards that could freely move in the sagittal plane. As hypothesized, we found a larger RMS of CoM accelerations in children and older adults and more balance losses in older adults compared to younger adults, indicating worse balance performance. Additionally, across age groups and conditions, the contribution of the CoP mechanism to the total CoM acceleration was dominant, i.e., 95%-108%, in accordance with our hypothesis. The contribution of the counter-rotation mechanism ranged between 19%-31% (with totals when summing the contribution of CoP mechanism and counter rotation mechanism higher than 100% indicating opposite effects of both mechanisms). Contrary to our hypothesis, only children and not the older adults did use the counter-rotation mechanism more to accelerate the CoM than younger adults.

### 4.1. Effects of surface instability

A total of 23 balance losses occurred spread over trials from three children, one younger adult and four older adults, all when standing on a balance board. The RMS of CoM accelerations was larger during standing on a balance board than on the rigid surface, reflecting that standing on the balance board was indeed more challenging than standing on the floor. Decreased pertinence of proprioceptive information from ankle muscles and a reduction of the effectiveness of ankle moments to accelerate the CoM when standing on a balance board could be an explanation for this (Horak et al., 2001; van Dieen et al., 2015). The number of balance losses (23), with balance boards that could freely move in the sagittal plane, was much larger than in our previous study (van den Bogaart et al., 2022), with uniaxial balance boards that could freely move in the frontal plane (3). Surface instability in the sagittal plane thus seems more challenging compared to surface instability in the frontal plane. This could be due to the fact that establishing CoP shifts by loading and unloading the legs by extensor and flexor muscle activity respectively, is possible in the frontal plane (Winter et al., 1993), next to applying ankle moments. Increasing the height of the balance boards did lead to larger deviations from the mean balance board orientation, increased frequency of balance board rotations and increased RMS values and relative contribution of the counter-rotation mechanism. In contrast, no effects of balance board height were found on balance boards that could freely move in the frontal plane (van den Bogaart et al., 2022).

### 4.2. Age effects

#### 4.2.1. Performance and kinematics

Older adults did lose balance more often than younger adults, indicating worse performance in older adults. Despite being healthy and non-falling, the older participants lost balance more often than the younger adult participants, which could indicate a higher risk of falls in daily life. Sensorimotor control is worse in children and older adults compared to younger adults, hence an increased RMS of CoM accelerations, corresponding to a deterioration of balance performance, and larger deviations from the mean board orientation compared to younger adults were expected (Bugnariu et al., 2006; Hirabayashi et al., 1995; Shams et al., 2020; Steindl et al., 2006; Teasdale et al., 1991). The differences in postural control between older adults and children on one hand and younger adults on the other hand during standing on a rigid surface or unstable surfaces are in line with previous studies (Bergamin et al., 2014; Hsu et al., 2009; Riach et al., 1989; Sturnieks et al., 2011). Despite the instruction to look at a marked spot on the wall in front of them at eye level, children did rotate their head more compared to younger adults. More head rotation in children could be self-generated and indicate less attention, which is common in children compared to younger adults (Huang-Pollock et al., 2002; Wickens, 1974). Self-generated head rotation may be disadvantageous, as it leads to changing visual and vestibular inputs, requiring more processing to discern between body motion and self-imposed head motion (Khan et al., 2013). Furthermore, self-generated head rotation could potentially result in increased muscle tone (e.g., leg and arm muscles) due to the tonic neck reflex (Bruijn et al., 2013; Iles et al., 1992; Parr et al., 1974). However, the head rotation in children was only around three degrees.

The higher frequency of balance board rotations in children and older adults could potentially reflect an increased frequency of CoM accelerations due to the CoP mechanism, and did coincide with an increased frequency of total CoM accelerations. However, increased frequency of the CoM accelerations did not lead to improved balance performance. It should be kept in mind that board rotations can reflect corrective actions as well as perturbations due to neuromuscular noise. Furthermore, the magnitude of CoM acceleration can indicate control of the CoM relative to the BoS, but perturbation effects on the CoM accelerations cannot be distinguished from control actions.

#### 4.2.2. Postural control mechanisms

The contribution of the CoP mechanism to CoM acceleration was dominant relative to the contribution of the counter-rotation mechanism. The relative contribution of the CoP mechanism to the total CoM acceleration was around 100% (ranging from 95%-108%) and the relative contribution of the counter-rotation mechanism was around 25% (ranging from 19%-31%). The contribution of the two mechanisms was not always in the same direction, as the summed RMS values were often larger than the RMS values of the total anteroposterior CoM acceleration. However, the desired direction for either of these mechanisms is unclear.

We found that children used the counter-rotation mechanism relatively more to accelerate the CoM compared to younger adults. This is in contrast to our previous study using balance boards that could freely move in the frontal plane, in which we did not find an effect of age on the relative use of the counter-rotation mechanism (van den Bogaart et al., 2022). The increased contribution of the counter-rotation mechanism in children cannot be explained by differences in body height between children and younger adults, as accelerating the body center of mass by the counter-rotation mechanism is less efficient at lower height (A.1.1. Supplementary materials). The increased amount of head rotation in children is unlikely to have contributed substantially to the increased rate of change of angular momentum (i.e., use of the counter-rotation mechanism) as head rotation was limited to only three degrees. We suggest that children are still learning to limit the contribution of the counter-rotation mechanism to the same extent as younger and older adults (Shumway-Cook et al., 1985). Overall, the contribution of the counter-rotation mechanism was limited, also in children. It could be that segmental rotations were used to achieve a proper orientation of segments such as regulating the orientation of the head in space, rather than accelerating the CoM (Alizadehsaravi et al., 2021). All participants, even the children, kept their head quite stable. This suggests that people prefer to maintain a constant visual and vestibular input by keeping the head stable, rather than using upper body rotations as a counter-rotation mechanism to accelerate the CoM. In addition, rotational accelerations of body parts need to be reversed leading to the opposite effect on the acceleration of the CoM. We also found limited use of the counter-rotation mechanism to accelerate the CoM in gait, as using the counter-rotation mechanism would actually interfere with the gait pattern (van den Bogaart et al., 2020). During unipedal stance on a balance board, larger contributions of counter-rotation were found, but this was to a large extent generated by the free leg (van Dieen et al., 2015).

## 5. Conclusion

Children and older adults had a poorer balance performance, than younger adults. Across age groups and conditions, the contribution of the CoP mechanism to the total CoM acceleration was much larger than that of the counter-rotation mechanism, ranging from 95%-108% vs 19%-31%. The CoP mechanism was dominant, likely because segmental rotations (i.e., the counter-rotation mechanism) were used to achieve a proper orientation of segments, such as regulating the orientation of the head in space, rather than accelerating the CoM. However, increasing the height of the balance board provoked increased use of the counter-rotation mechanism. Furthermore, children did, but older adults did not, use the counter-rotation mechanism relatively more to accelerate the CoM compared to younger adults, probably due to the fact that children are still learning to limit the contribution of the counter-rotation mechanism to the same extent as younger and older adults.

## 6. Acknowledgements

The authors gratefully acknowledge The Expertise centre for Digital Media (EDM) for technical support and Kimmy Daenen and Charlotte Uwents for their help during the experiments.

## A.1. Supplementary materials

### A.1.1. The effect of body height on the contribution of the counter-rotation mechanism to CoM acceleration

As described in the manuscript, the contribution of changes in angular momentum to whole body acceleration is;

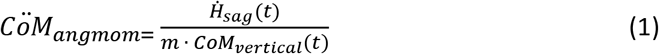

Now, let us assume;

1. that the angular velocities that are involved are similar between age groups
2. that changes in angular momentum are proportional to angular momentum itself (i.e. 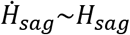
3. that mass scales with body length squared (i.e. *m*~*l*^2^)

angular momentum for the whole body in the sagittal plane can be described as:

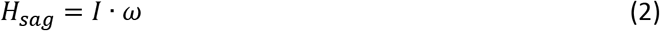

With ω being the total body angular velocity (see assumption 1) and *I* being the inertia tensor for the combined body, which can be calculated as the sum of the segment masses times the square of their moment arm with respect to the CoM:

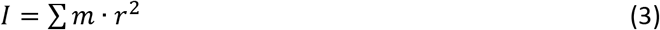

Where *m* is the mass of a segment, and *r* is the moment arm of the segments CoM with respect to the full body CoM. Now, since mass scales with body length squared (assumption 3), and *r* also scales with body length, equation 2 can be rewritten as:

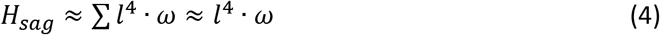

Using assumption 2, and putting (4) into (1) (and substituting the *m* in the denominator, as well as the *CoM_vertical_*(*t*), which is also proportional to *l*, we get;

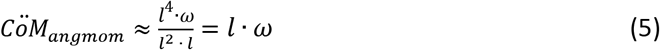

Thus, the contribution of changes in angular momentum to whole body acceleration increases linearly with length.

